# Markov chains applied to molecular evolution simulation

**DOI:** 10.1101/653972

**Authors:** Joel Valdivia Ortega

## Abstract

Today we are able to describe genome evolution but, there are still several open questions on this area such as: Which is the probability that a mutation occurs at a nucleotide level? Is it possible to predict the evolution of a particular genome?, or talking about preservation, is there a way to simulate the genetic diversity for endangered species? In this paper it is shown that it is possible to make a mathematical model not only of mutations on the genome of species, but of evolution itself, including factors such as artificial and natural selection. It is also presented the algorithm to obtain the probabilities of mutation for each specific part of the genome and for each specie. The potential of having this tool is giantic going from genetic engineering applied to medicine to filling up blank spaces in phylogenetic studies or preservation of endangered species due to genetic diversity.

## Main text

Nowadays the work done in the genetics and molecular evolution field is based on describing and manipulating genomes *(1,2),* while the few works on making a model that allow us to make predictions on the evolution of genome in species, and therefore understand the phenomena more clearly, are all based on suppositions over the probability distribution the mutations follows *(3)*.

In this paper, no probability distribution will be assumed, but we will obtain the probabilities of transition just as a result of analysing the genomes of linked species. In order to do this we will use Markov chains, which is a mathematical method used to describe the evolution of discrete systems through time. By having this, we will then be able not only to simulate the most probable evolution between species but we will also demonstrate that it is possible to rebuild one genome based on the predecessors or vice versa.

To explain the way Markov chains work, let *N* be the set of possible states our system can take, *x*_*n*_ be a random variable at the n-th moment which takes values on *N* and *P* a matrix called “transition matrix” such that the entry *P* _*i,j*_ is the probability that *x*_*i*_ is in the i-th state and that *x*_*n*+1_ is in the j-th state. Therefore we have λ_*n*+*m*_ = *P*^*m*^*x*_*n*_, where λ_*n*+*m*_ is the probability distribution of *x*_*n*+*m*_, noting that *m* can also be negative.

The resemblance between this and the molecular evolution is already tangible, so let us take *N* = {*A, C, G, T*} as the ordered set of nucleotides, where *A* stands for adenine, *C* for cytosine, *G* for guanine and *T* for thiamine.

In general, the next step would be choosing at least two linked species, where by linked we mean that one is the ancestor (or sucesor) of the other(s), as some of the known values our random variable must take, then we obtain the matrix *P*^*m*^ where *m* is the number of generations between the selected species and as soon as we calculate its *m* -th root, we are finished because once we have *P*, we are able to study the whole genomic history of our species, both future and previous generations.

To give an example and for computational reasons, we will take *x*_0_ as the first ten thousand nucleotides of the wolf’s genome according to the National Center for Biotechnology Information (NCBI) *(4)* and *x*_*n*_ as the first ten thousand nucleotides of dogs such as a boxer, a poodle, a beagle and a yorkshire terrier *(5-8)* from the same source.

According to The Smithsonian *(9),* the domestication of dogs occured around 30,000 years ago so assuming each specimen had at least 1 son per year, let us suppose that there are around 30,000 generations between our wolf and our dogs and that the transition matrix is the same for all of them, giving us more information so our probabilities might be more precise.

To obtain *P* ^30 000^ it is only needed the combinatorial probability, which consist on organizing the genomes in vectors, counting the amount of times the *i* -th nucleotide of the wolf remained the same or changed to another one for the four dogs and for each entry of our vector and then normalizing the matrix such that the sum of the entries of all rows is equal to 1 in order to understand the transition matrix as a matrix of probabilities. In our example, the result is

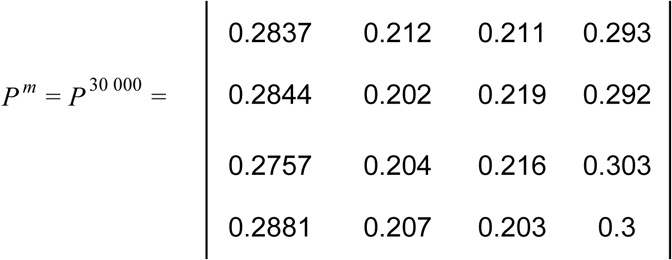

The problem with remaining with this matrix is that we would only be capable of analysing the molecular evolution making jumps of 30 000 generations each time, not to mention the that this matrix do not tell us much information since every row corresponds to an almost uniform distribution between the 4 nucleotides.

In order to achieve our goal, we will define a n-th root to our matrix. To do that we will calculate the eigenvectors of *P*^*m*^, place them as the column vectors of a matrix *B* and obtaining the inverse matrix of *B, B*^−1^. Then we calculate *D*^*m*^ = *BP*^*m*^*B*^−1^, which will be a diagonal matrix with the eigenvalues of *P*^*m*^ as the non zero entries, and apply the n-th root to the numbers in the diagonal so we get *D*. After all this, *P* will be the matrix of the real entries of *B*^−1^*DB*, being worthy to say that the imaginary part of P may appear because the eigenvalues of *P* ^*m*^ may be complex numbers. With this process we get our transition matrix, which for our example is

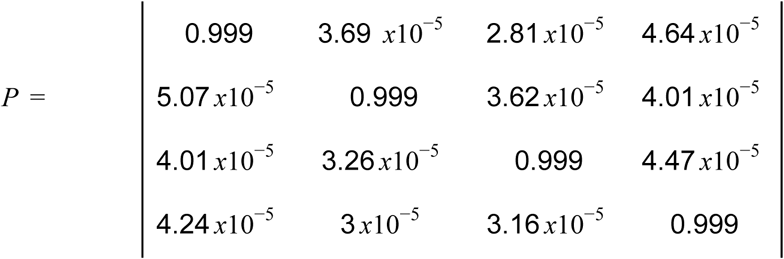

Now we have our transition matrix we are able to do some experiment with it in order to see if Markov chains are a good model for evolution, but let us first analyze *P* and see if it has correspondance with what we see in our daily experience.

Everytime we see the puppies of some dog, we can appreciate the resemblance between parents and sons, so this means that our matrix should not make too many mutations from one generation to a successive one. On the other hand we know that evolution happens, so our matrix should leave space for mutations to happen from one time to another.

This is accurately represented on the entries of *P* because it says that the probability that a nucleotide remains the same is almost one but still leaves some probability for mutations to occur because probabilities of change are not null, so at least at a mathematical level, our transition matrix makes sense.

But before we go on, we might make just some more observations about *P* and with those get some mathematical explanations of the way evolution work, to show the power of having our transition matrix. For example the fact that no single entry of the transition matrix is zero, tells us it is not possible to separate evolution into several Markov chains; the fact that there are any attractor subset, tell us that evolution will not get stuck nor finish at any point; since no column vector is null, it is not likely to happen that someday one nucleotide will be totally replaced by the other ones; and since the probabilities of change for each row are very similar between each other, we can affirm that molecular evolution has no prefered direction and that it works as a *random walker* between nucleotides.

Since up to this point things have gone as we expected, we are now interested in testing our model with the most basic probe it should accomplish in order to be a good model of evolution: beginning with the wolf genome and having the transition matrix, a computational program should be able to obtain the genome of the four dogs we are considering.

In order to do this we wrote down a *artificial selection* program which, beginning with the genome of the wolf and choosing a specific dog as a target, simulated 50 children and chose the 2 of them which were more similar to the dog. After doing that, the program simulated the chosen children to have a breed between themselves and the wolf and chose the specimen which had the closest genome to the dog we were attempting to reach, simulating now that it had 50 children. This process was applied iteratively expecting that at some point the program will converge.

The process was made for all the four dogs and in every case the program converged but since this is a probabilistic method, now we must place the question, does it has to be with the dogs or we are just reaching the desired genome for the way the algorithm was made?

To answer this, we chose a randomly generated genome as the target and ran the program in order to see if the program also converges in this case and if so, if there was a qualitative or quantitative difference between the other cases. The results of simulating 100 times the process are shown below

As we can see on Table1, there is no significant difference between the results from the dogs and the results from the randomly generated genome so this means that this *artificial selection* program is not good to predict the genome’s evolution outside the known species.

**Table 1:** Statistical data about the results of computational process to reach to the dogs genome and the randomly generated genome

|  | Boxer | Poodle | Beagle | Yorkshire terrier | Random genome |
| --- | --- | --- | --- | --- | --- |
| Average amount of generations taken | 1036.434 | 996.94 | 1042.538 | 1062.708 | 1077.428 |
| Standard deviation | 135.023 | 127.864 | 137.112 | 135.142 | 134.419 |
| Standard deviation as a percentage of the average | 13.027 | 12.825 | 13.151 | 12.716 | 12.475 |

But not everything is lost because the fact that the program drive us with probabilistic methods from the wolf’s genome to each of the dog’s genome tells us that this *artificial selection* program give us the most probable molecular evolution between the known species, letting us know the “genomic history” that connects one specie with the other.

But we are trying to design a way to study the whole process of evolution, not just a interpolation between species so what we made was, using our transition matrix and for computational simplicity selecting just the first *n* nucleotides from each dog’s genome, to simulate 7 probable parents for each of our 4 dogs and then to choose the closest ones between each other. After that, we iterated the simulation for the selected parents and ran the program until it converges with a difference between the parents of 1%.

Once we arrived to this difference, we calculated the percentage of differences between the genome we arrived to and the wolf’s genome. Given a *n*, we repeated this process 20 times for statistical purposes and we called *discordance of n (D*_*n*_ ) to the average this quantities.

On the other hand and keeping the given *n*, we calculated the average percentage of similarity between the chain of nucleotides of each dog and the one from the wolf and to that number we called *concordance of n (C*_*n*_ ).

It is worthy to remark that, since we did not use directly the wolf’s genome in this algorithm but to get the transition matrix, whatever the result is, it will not depend on anything but the genomes from the beginning and the transition matrix.

We took 74 different *n* ‘s represented on Figure 1 and the result was that the average sum of the concordance and the discordance of each *n* (*C*_*n*_ + *D*_*n*_) is 100.03% with a standard deviation of 2.81%, allowing us to take this as a statistical truth for all *n*

**Figure 1:**
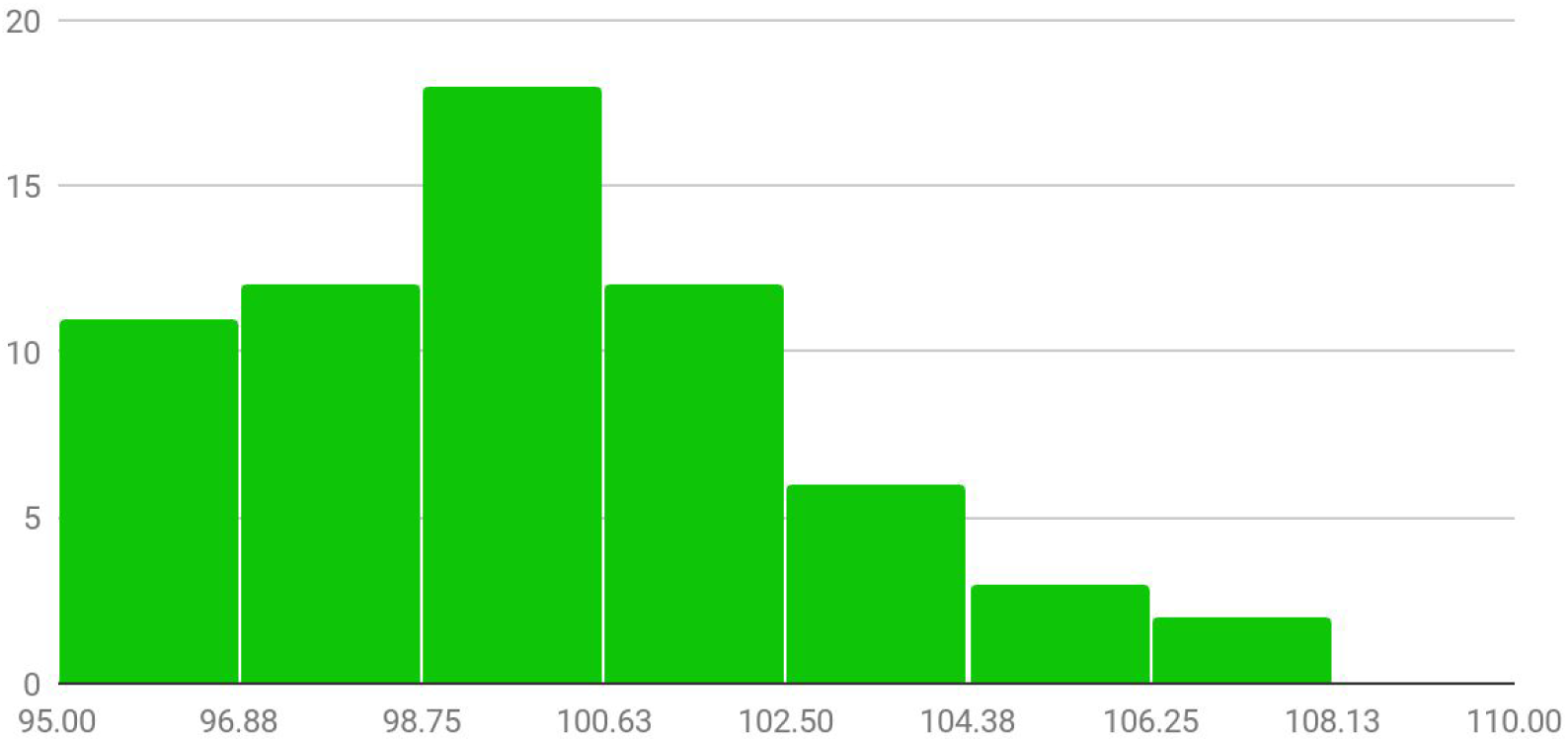
Distribution of discordance puls concordance for several n.

This result is the final confirmation that modelling evolution as a Markov chain is accurate, that our transition matrix has the correct probabilities of transition between nucleotides and that we cannot only interpolate but also extrapolate predictions of the genome’s evolution through given species because we know that as bigger the chain of DNA we take, the more similar the dog’s genome will be to the wolf’s one and therefore the discordance will tend to zero, i.e.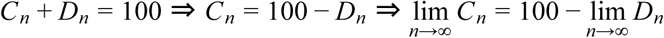, but from statistical biology we know that 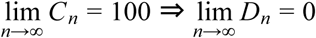.

In other words, the probability that the resulting genome with this algorithm converges to the wolf’s genome tends to 1 as we take more nucleotides in our genome, or written in mathematical symbols 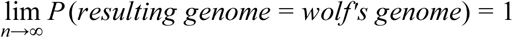.

It is remarkable the paper the amount of used dogs had in this work, because that is what gave us the extra information so the program was able to converge to a similar genome to the wolf’s one.

